# Genetic and real-world clinical data, combined with empirical validation, nominate JAK-STAT signalling as a target for Alzheimer’s Disease therapeutic development

**DOI:** 10.1101/179267

**Authors:** Alejo J. Nevado-Holgado, Elena Ribe, Laura Thei, Laura Furlong, Miguel Angel-Mayer, Jie Quan, Jill C. Richardson, Jonathan Cavanagh, NIMA Consortium, Simon Lovestone

## Abstract

As Genome Wide Association Studies (GWAS) have grown in size, the number of genetic variants that have been nominated for an increasing number of diseases has correspondingly increased. Despite this increase in the number of associated SNPs per disease, their biological interpretation has in many cases remained elusive. To address this, we have combined GWAS results with an orthogonal source of evidence, namely real-world, routinely collected clinical data from more than 6 million patients in order to drive target nomination. First we show that when examined at a pathway level, analysis of all GWAS studies groups Alzheimer’s disease (AD) in a cluster with disorders of immunity and inflammation. Using clinical data we show that the degree of comorbidity of these diseases with AD correlates with the strength of their genetic association with molecular participants in the JAK-STAT pathway. Using four independent open-science datasets we then find evidence for altered regulation of JAK-STAT pathway genes in AD. Finally, we use both in vitro and in vivo rodent models to demonstrate that Aβ induces gene expression of key drivers of this pathway, providing experimental evidence validating these data-driven observations. These results therefore nominate JAK-STAT anomalies as a prominent aetiopathological event in AD and hence potential target for therapeutic development, and moreover demonstrate a de-novo multi-modal approach to derive information from rapidly increasing genomic datasets.

**One Sentence Summary:** Combining evidence from genome wide association studies, real-world clinical and cohort molecular data together with experimental studies in rodent model systems nominates JAK-STAT signaling as an aetiopathological event in Alzheimer’s disease

## Introduction

As Genome Wide Association Studies (GWAS) have grown in size, often now numbering tens of thousands of research participants, the numbers of genes with common variants that contribute to disease susceptibility have correspondingly grown. This is as true for Alzheimer’s Disease (AD) as it is for many other disorders, and bioinformatic and pathway analyses of these large number of susceptibility genes is providing a highly efficient method of proposing and prioritising underlying biological pathways of these disorders for further study *(1, 2)*. In some cases this understanding adds to existing observations – such as the evidence from GWAS that pathways of inflammation are important in AD – whereas in other cases pathway analysis yields less expected findings, such as evidence that cholesterol synthesis and endocytic recycling are part of the pathological process *(3)*. However, such analyses have their limitations, not least due to the fact that our understanding of molecular pathways is far from complete. Thus, many pathways described as categorical entities through data sources such as Kyoto Encyclopedia of Genes and Genomes (KEGG) and similar approaches, are skeletal at best and only contain a fraction of the genes that might be involved in any given process. Furthermore, many genes, including hub genes of networks but also others, are nominated in multiple different pathways through their pleiotropic function. As an example, one of the most replicated associations with AD, the gene CLU encoding clusterin, is involved in processes as diverse as complement signalling, protein binding and chaperoning and cell survival *(4)*. In the context of this incomplete understanding together with known and unknown molecular pathway complexity, determining the underlying biology of disease from GWAS studies becomes difficult and hence inevitably limited.

In an effort to address this limitation, we reasoned that it should be possible to hone pathway analysis utilising orthogonal datasets. Specifically, we hypothesised that shared molecular pathways might be more meaningful to an understanding of the molecular processes of aetiopathogenesis if diseases that shared pathways also shared morbidity. Put another way, if two or more diseases are more commonly found to co-occur rather than by chance, and if those comorbid diseases also share molecular pathways, one would predict that those shared pathways are more likely to play a role in aetiopathogenesis. In order to test this reasoning, we combined pathway analysis of the GWAS associations from all diseases (as reported in the GWAS catalog; https://www.ebi.ac.uk/gwas/) together with a co-morbidity study from real-world data to identify shared pathological processes. We then tested the resulting pathway in observational and empirically derived genome wide expression datasets from human and rodent studies, and finally validated the results in empirical studies in rat models in vitro and in vivo. The results, demonstrating a role for JAK-STAT signalling in AD, are in line with the known contribution of inflammatory processes to the disease, but they further nominate a specific target for therapy and provide a possible approach to interpretation of GWAS data for other disease areas.

## Results

### Subhead 1: Genes associated with AD show shared susceptibility to diseases of immunity

Given that recent large scale association studies suggest genes related to AD risk are involved in many different biological pathways, only some of which would have been predicted in advance, we wondered whether some of these pathway associations would be shared with other diseases. To address this question systematically, we first obtained from the GWAS catalogue *(5)* a list of all genes that have been found to be associated with all diseases in that dataset. Subsequently, for each gene appearing in the list, we determined the number of KEGG pathways that that gene shared with each other gene on the list. As a general concept, we assume at this point that if two genes, one associated with one disorder and the other associated with a second disorder, both appear in the same biological pathways, then those pathways should commonly be associated with both diseases.

Using this data of genes associated with diseases and the KEGG pathways that those genes appear in, we then generated a gene-gene matrix, plotting for each gene associated with each disease the numbers of KEGG pathways shared with genes associated with each other disease in the dataset. Not surprisingly, we find many genes participate in many different pathways. This is illustrated for three disorders in Fig. 1, using for this explanatory purpose five genes only. In this illustrative segment of the data, it can be seen that the genes associated with Crohn’s disease in particular participate in many different named KEGG pathways. Again, this is unsurprising as these genes are part of the Human Leukocyte Antigen system encoding the Major Histocompatability Complex; a much studied master regulator of immunity. Using this gene-gene matrix we are then able to explore which diseases share pathways with other diseases.

**Fig. 1.**
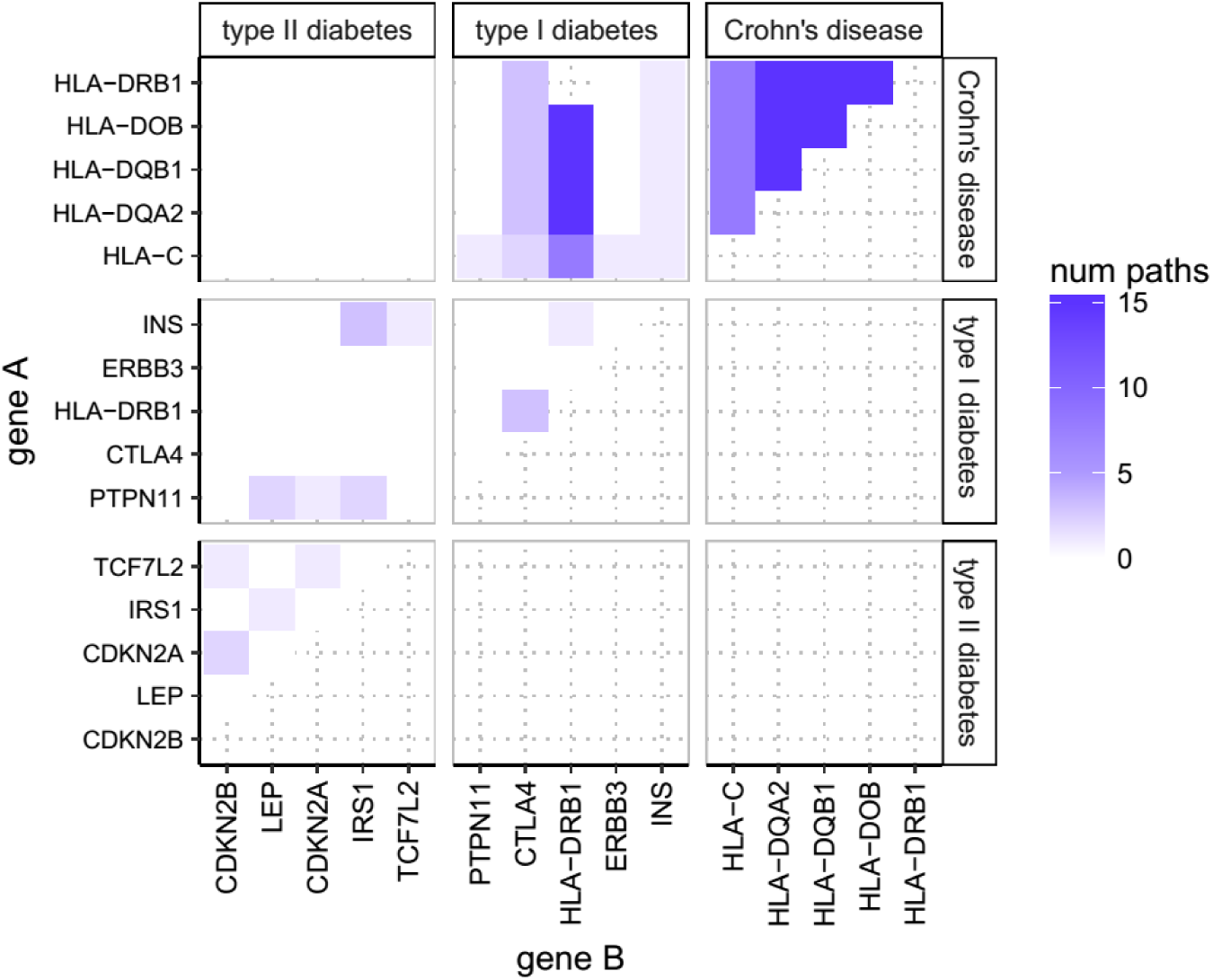
Number of shared pathways. Number of pathways shared by the top 5 genes of 3 of the studied diseases - Crohn’s Disease and Type 1 and Type 2 Diabetes Mellitus. For any two genes (as indicated on the X and Y axes), the color of the corresponding cell represents how many KEGG pathways these two genes share. Each one of these 15 genes has been associated through a GWAS study with at least one of these diseases. Although performed for each disease in the GWAS Catalogue using the 20 most strongly associated genes (i.e. lowest p-value), this figure only shows the top 5 genes and only 3 diseases for representational purposes.

Again in the illustrative segment of the data in Fig 1 it can be seen that the genes associated with Crohn’s disease share no pathways with the genes associated with Type II diabetes (T2DM), but do share considerable numbers of KEGG pathways with genes associated with Type 1 diabetes (T1DM). Note that this observation is not driven by the same genes being associated with any two diseases, as we exclude from the comparison the number of pathways that HLA-DRB1 shares with itself in the T1DM-Crohn’s instance. Rather the T1DM-Chrohn’s association is driven by different genes sharing common pathways such as CTLA4 and INS, or HLA-DRB1 with genes others than itself, which are genes of the HLA complex genes associated with both T1DM and Crohn’s disease.

The GWAS catalogue includes 59 diseases where studies have identified at least 25 associated genes per condition, and these were therefore used in this analysis. Using the Wilcoxon rank-sum test, most of these conditions did not show significant overlap of pathways derived from GWAS genes in the pathway space, except for a cluster of 18 disorders that strongly overlapped with each other (Bonferroni corrected p-value < 0.05). 14 of these diseases were disorders commonly classified as diseases of immunity, while the other 4 were AD, Age Related Macular Degeneration (ARMD), T1DM and Hypothyroidism (see Fig. 2 for a segment of the data and Fig. S1 for the full matrix). Part of this cluster is evident in Fig. 2 as 11 diseases that overlap with each other in 44 out of 55 disease-disease pairs (repeated pairs eliminated) while the full cluster of 18 disorders includes 127 out of 153 significant pairs as seen in Fig. S1. The disorder with fewest associations with other diseases from the cluster was Seasonal Allergic Rhinitis (8 out of 18), while the disorders out of the cluster which had highest number of associations with the cluster were Atopic Eczema and HIV-1 Infection (both 2 out of 18).

**Fig. 2.**
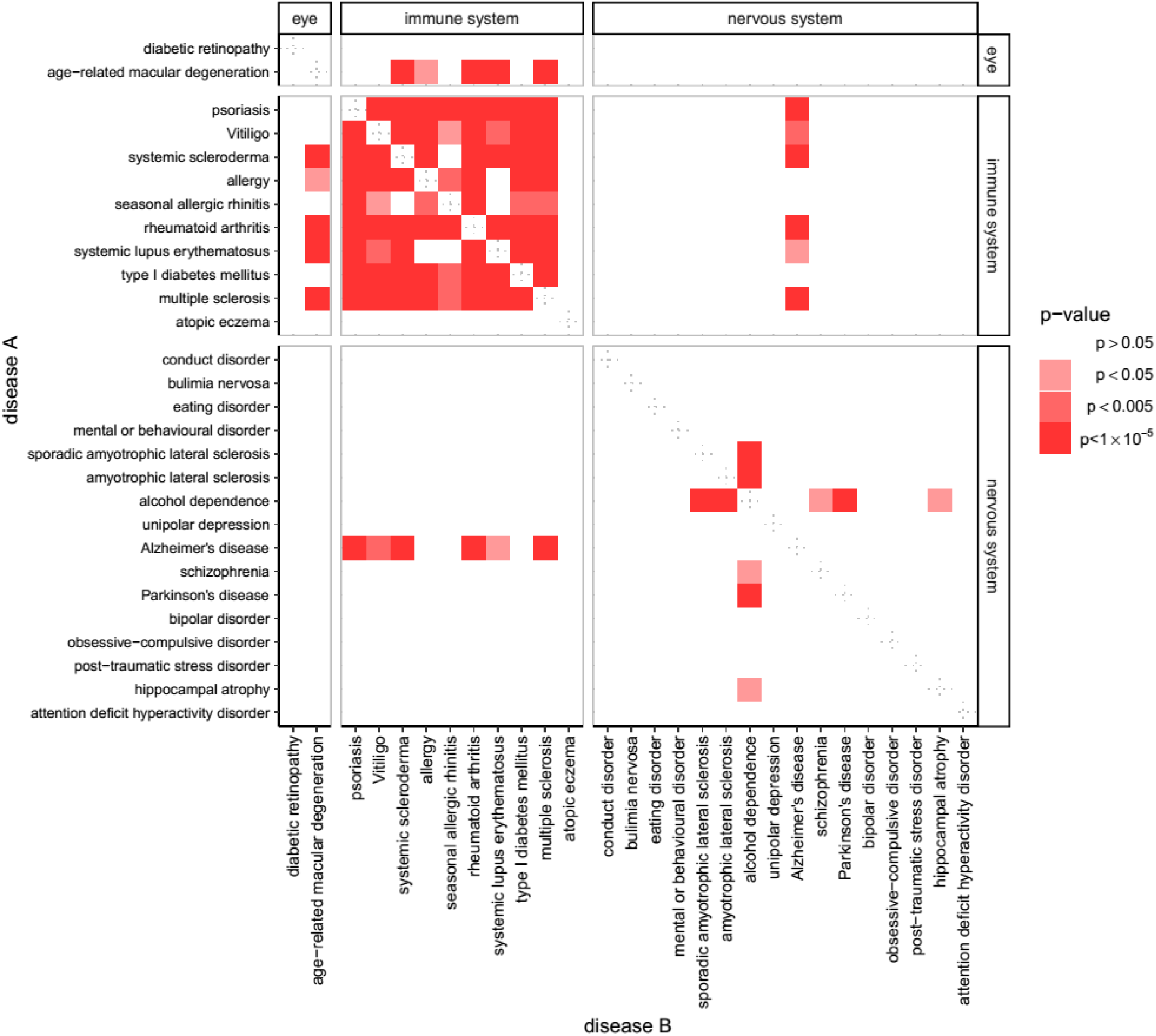
Overlap of molecular pathways in susceptibility genes for all disease. Susceptibility genes from association studies of multiple diseases overlap in the pathway space. Although we analyzed 59 diseases with at least 25 susceptibility genes each, for representational purposes the figure shows those classified as either eye disease, immune system disease, or nervous system disease. The color of each square in the table represents the p-value obtained when calculating whether the GWAS-genes of diseases A and B overlap in the pathway space more than would be expected from chance. Full data is presented in supplementary Fig. S1.

### Subhead 2: Among all KEGG pathways, associations with JAK-STAT signaling are shared between diseases co-morbid with AD

Having demonstrated that AD, together with ARMD, hypothyroidism and T1DM, share multiple KEGG pathways with disorders of immunity, we then explored which of these pathways was responsible for this overlap. In order to do this systematically, we first determined the strength of association of each disease with each KEGG pathway, calculating the proportion of susceptibility genes that a given disease has for each KEGG pathway (‘pathway load’). Then, using the US National Hospital Discharge Survey (NHDS), a dataset including diagnostic information from more than 6 million patients, we calculated the co-morbidity between disorders of immunity and late onset AD on discharge from in-patient care between 1979 and 2006. Finally, and for each individual pathway, we then calculated the degree of correlation between pathway load and comorbidity with AD. Most of the KEGG pathways showed no significant correlation, with only three exceptions, indicating possible meaningful shared pathways contributing to disease comorbidity (see Fig. 3). Of these, ‘Cytokine-cytokine interaction’ (KEGG ID hsa04060) and ‘Viral myocarditis’ (hsa05416) did not survive Bonferroni correction for multiple comparisons (uncorrected p value 0.05), while ‘JAK-STAT’ (hsa04630) did survive (uncorrected p value 0.00022) suggesting that the shared GWAS genes on this pathway might contribute to the observed co-morbidity between AD and these disorders of immunity

**Fig. 3.**
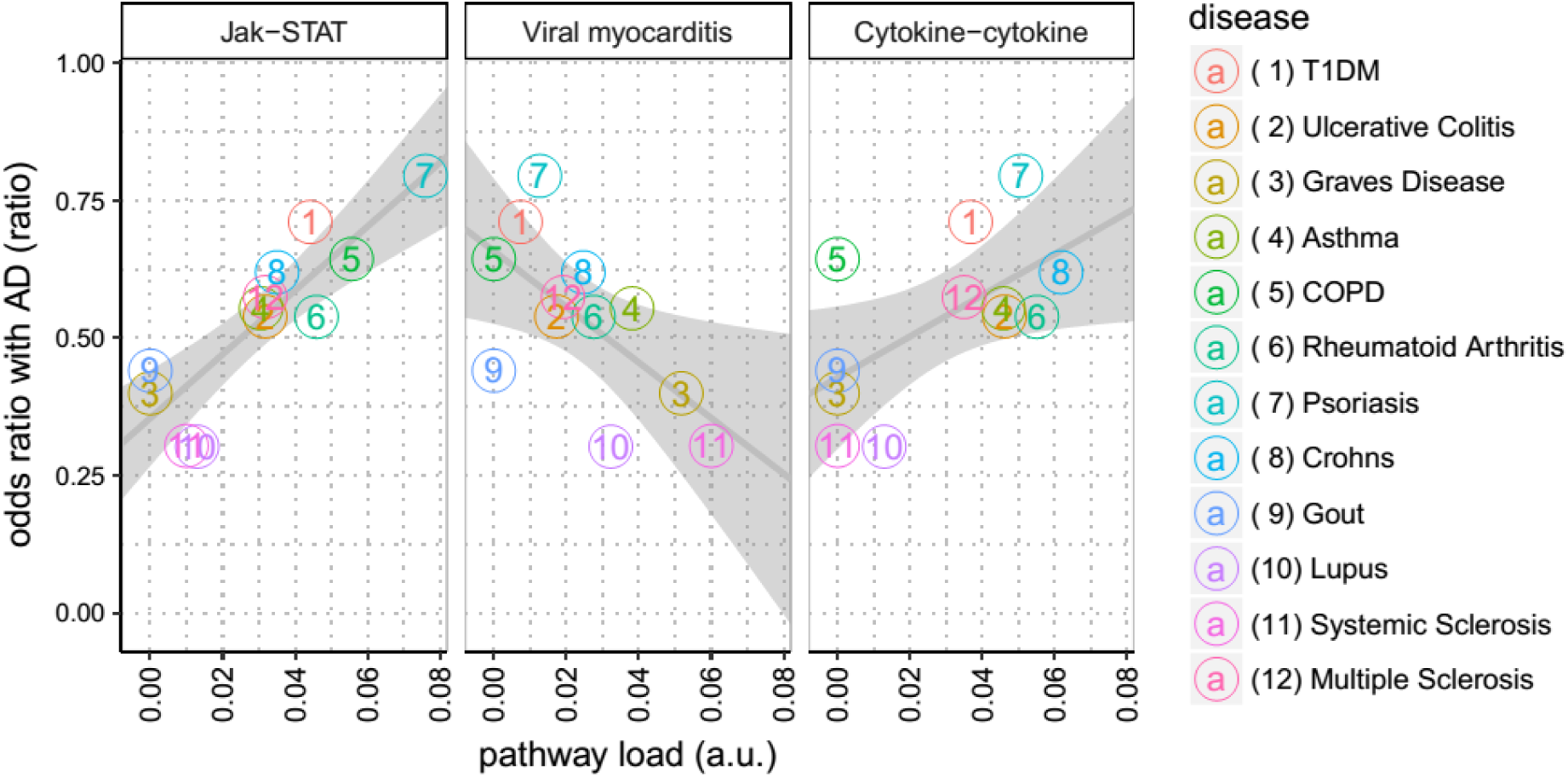
Pathways association with disease from real-world data. As described in the methods, we calculate the correlation between the co-incidence of the inflammatory diseases (denoted ratioAD(A)) and the proportion of susceptibility genes of these diseases on any given pathway (denoted load(A,p)).This figure shows this correlation for the three pathways where such correlation was the most significant. The ratioAD(A) of each disease is represented in the Y-axis, while load(A,p) is represented in the X-axis. Each disease is represented by a number and a color, as shown in the legend to the right. The p-values corresponding to each pathway are, from left to right, 0.00022, 0.027 & 0.039. Only JAK-STAT signaling survives multiple testing correction. T1DM stands for Type 1 Diabetes Mellitus; COPD for Chronic obstructive pulmonary disease; Cytokine-cytokine for Cytokine-cytokine receptor interaction pathway; JAK-STAT for JAK-STAT signaling pathway.

### Subhead 3: Evidence for altered JAK-STAT pathway gene expression in AD blood, brain and in an in vitro model with established relevance to AD

If the KEGG pathway is responsible for the observed co-morbidity between AD and disorders of immunity, then we reasoned that altered JAK-STAT signalling would be a feature of AD. In order to explore this, we utilised gene expression datasets from blood from human cohort studies (Datasets 1 and 2), from brain from post-mortem human studies (Dataset 3) and from a rodent in vitro model relevant to AD (Dataset 4) with robust proof of concept using empirical studies with gene-knockdown, in animal models and in post-mortem human brain.

We first examined expression of genes from the JAK-STAT pathway (KEGG ID hsa04630) in blood from patients with AD compared to unaffected controls, reasoning that post-mortem brain might have more late-stage, secondary changes of inflammation, possibly less relevant to aetiopathological pathways. We used two cohorts, one a multinational longitudinal study (AddNeuroMed - ANM; 105 AD and 114 unaffected age matched subjects; denoted Dataset 1 in this manuscript), and another, a single centre longitudinal study utilising the exact same protocol (Dementia Case Register - DCR; 95 AD and 78 control blood samples *(6)*; denoted Dataset 2). In ANM, 15 of the 47 JAK-STAT genes measured in this dataset were significantly dysregulated when comparing controls with AD subjects (p<0.05 after Bonferroni correction, see Fig. 4A). A binomial exact test reveals that this proportion of significant genes in the JAK-STAT pathway is larger than expected by chance (p-value 5×10^-9^). In DCR, 6 of the 47 JAK-STAT genes were significantly altered (p<0.05), also revealing significance at the pathway level (p-value 0.03). P-values obtained in ANM were significantly correlated (p = 4×10^-7^ for Spearman correlation, see Fig. 4B) with those obtained from DCR. We also merged both datasets and repeated the analysis, finding again 15 out of 47 genes were dysregulated (p<0.05) in the AD population when compared with the control population. We then examined JAK-STAT genes in the MCI population compared to unaffected controls in both cohort datasets, finding 10 out of 47 genes dysregulated in both studies, with binomial test being also significant (p-value 9×10^-5^).

**Fig. 4.**
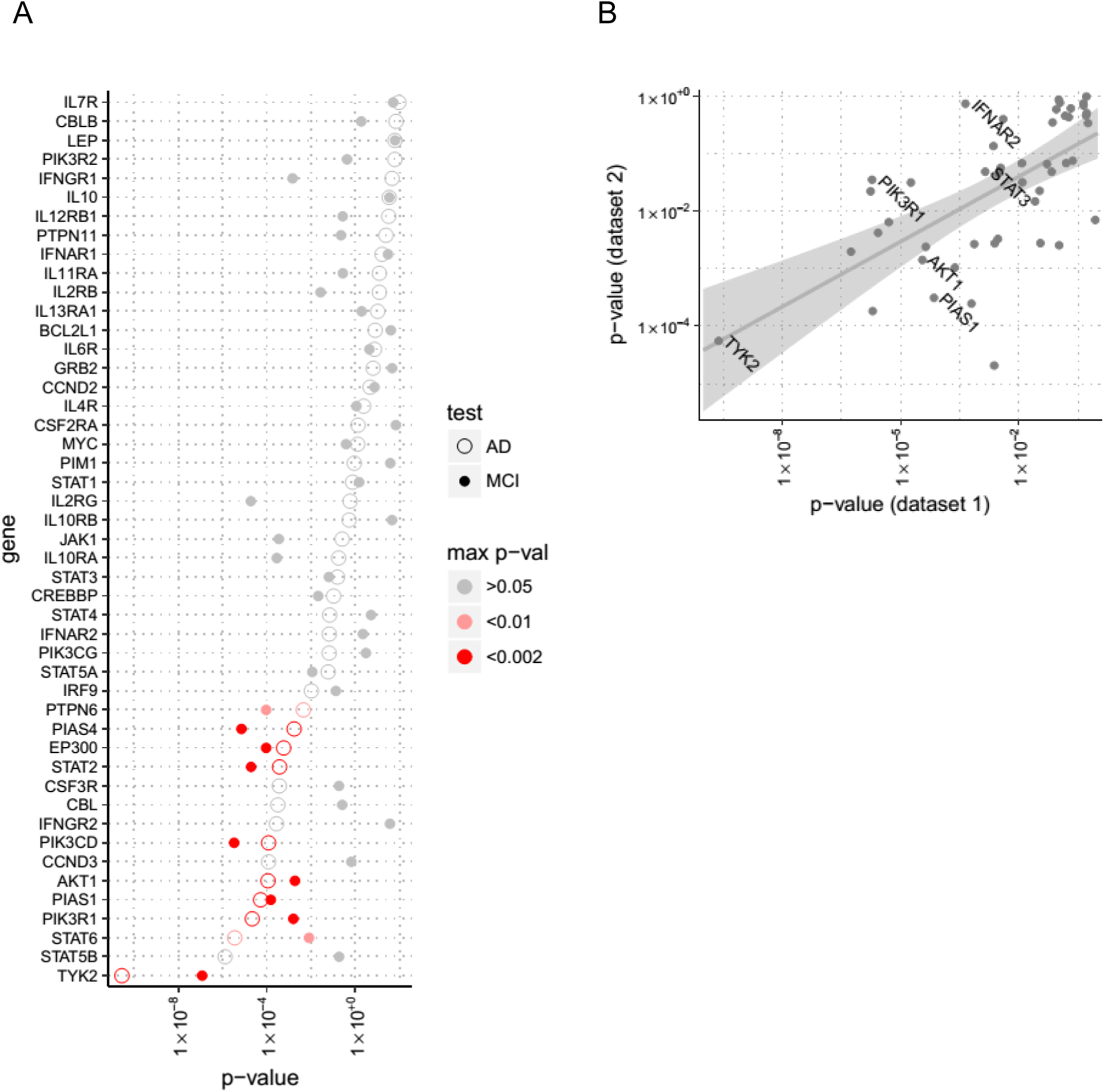
Gene expression of JAK-STAT pathway genes in blood in AD. (**A**) We determined the expression of JAK-STAT pathway genes in blood, comparing people with AD (open circles) and MCI (solid circles) to unaffected individuals in the ANM/DCR study. (**B**): Results from datasets 1 (ANM) and 2 (DCR). Axis represent p-values from each dataset. The solid line shows the a linear regression, with confidence intervals in grey shading. Names are shown for the most significant genes that were also sampled in dataset 3.

We then examined the expression of JAK-STAT signalling genes in post-mortem brain tissue utilising a study of 129 AD subjects and 101 controls *(7)* (Dataset 3). In this cohort, 153 JAK-STAT genes were sampled, of which 53 showed significant dysregulation (p-value < 0.05), and 37 of which survived multiple comparison correction (p-value < 0.05 after FDR adjustment). A binomial exact test reveals that this proportion of significant genes in the JAK-STAT pathway is larger than expected by chance (p-value 2×10^-16^). With respect to the five genes that were significant in ANM (Dataset 1) and that were also sampled in DCR (Dataset 2), TYK2 (p-value 0.0002), IFNAR2 (0.04) and PIAS1 (2×10^-5^) were also significantly altered in Dataset 3 (see Table 1).

**Table 1.** Comparing most significant genes P-values from analysis of case versus control samples when comparing RNA-expression in the different datasets. Among the 15 most significant genes of ANM (Dataset 1), the table shows those that were also sampled in DCR. (ANM-AddNeuroMed; DCR-dementia case register; AE-ArrayExpress – see methods for details)

|  | Dataset 1 | Dataset 2 | Dataset 3 | Dataset 4 |
| --- | --- | --- | --- | --- |
| Gen | (blood, ANM) | (blood, DCR) | (brain, AE) | (in-vitro) |
| TYK2 | $5 \cdot 10^{-10}$ | 0.0005 | 0.0002 | - |
| PIK3R1 | $1 \cdot 10^{-6}$ | 0.03 | 0.3 | 0.5 |
| IFNAR2 | 0.0004 | 0.7 | 0.04 | 0.007 |
| AKT1 | 0.0003 | 0.001 | 0.5 | 0.001 |
| PIAS1 | 0.0007 | 0.0003 | $8 \cdot 10^{-5}$ | 0.016 |

Finally, we also analyzed a previously reported *(8)* in vitro cell model derived dataset (Dataset 4). This study demonstrated that in rodent neurons, exposure to Aβ induced gene expression changes that overlap extensively with changes in gene expression induced by the Wnt signaling antagonist, Dickkopf-1 (Dkk1); itself a gene induced in response to Aβ in rodent cells, animal models and human brain. This gene expression signature was subsequently validated in animal models with cerebral amyloid plaque pathology and in AD brain, as well as in a human DKK1 over-expressing mouse model. Knock-down of particular genes on this pathway prevented Aβ induced phenotypes including neurotoxicity and tau-phosphorylation. This experimental dataset contained measurements of 32 JAK-STAT genes, with 8 of these being significantly dysregulated (p-values < 0.05). Binomial testing showed that this proportion of dysregulated genes was higher than expected by chance alone (p-value 0.0001). Of the four genes that were significant in ANM (Dataset 1) and were also sampled in all other datasets (DCR – Dataset 2, Array Express – Dataset 3; and rodent model – Dataset 4), Akt1 (p-value 0.001), Ifnar2 (0.007) and Pias1 (0.016) were also dysregulated in Dataset 4 (see Table 1).

### Subhead 4: Empirical evidence for JAK-STAT dysregulation in both in vitro and in vivo models of Aβ-induced neurotoxicity

We then further validated these bioinformatics results using empirical studies, measuring the key drivers of the pathway, Jak1, Jak2, Jak3 and Tyk2, in an in vitro model of neuronal toxicity.

Previously, we and many others have demonstrated that rodent neurons exposed to amyloid peptides are susceptible of Aβ-induced toxicity and other phenotypes, including tau phosphorylation and synaptic alterations *(8–12)*.

We therefore exposed primary neuronal cultures to 3 μM of oligomerised Aβ42 and real time PCR was performed at 30m, 4h and 24h after Aβ exposure (Fig. 5). Hprt1 and Actb were used as housekeeping genes for data normalization. Tyk2 showed a time-dependent increase in expression detectable at 4h (p=0.0077) and maintained up to 24h (p= 0.0063). Jak1 and Jak2 mRNA levels were only significantly increased at 24h with p values of p= 0.024 and p= 0.015, respectively. Jak3 expression was modulated by Aβ at any time point (p>0.05).

**Fig. 5.**
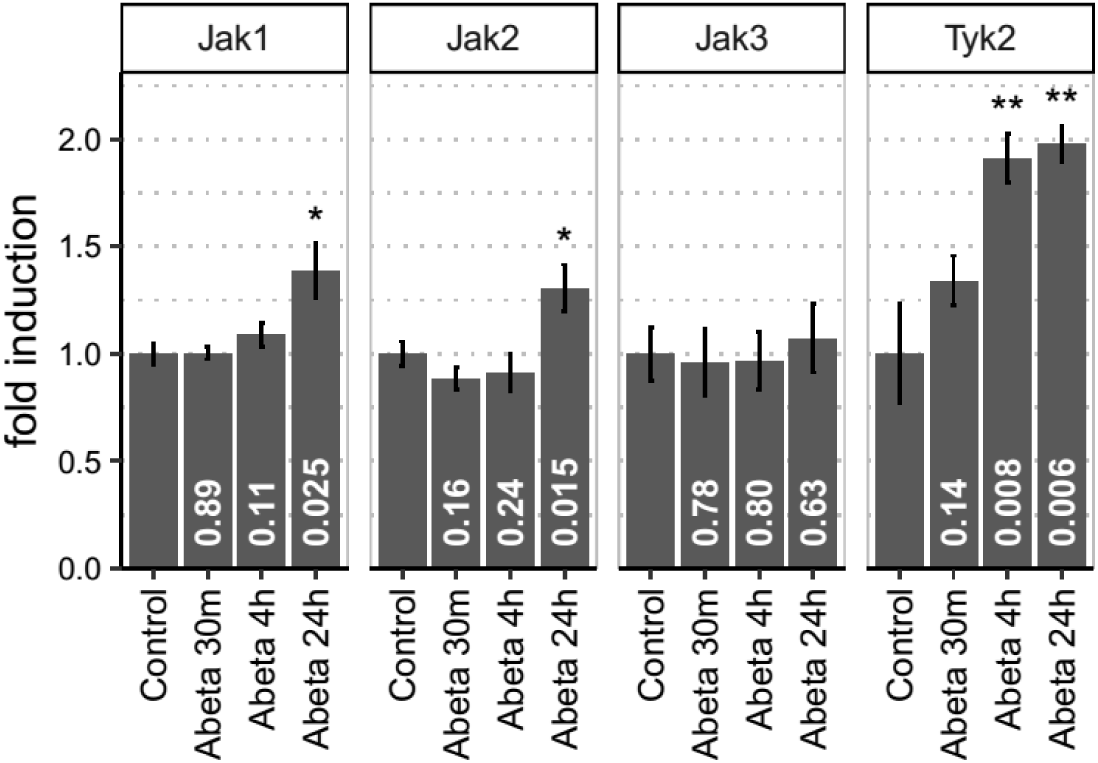
The Janus kinases Jak1, Jak2 and Tyk2 are induced in vitro after Aβ stimulation in a sequential pattern Neuronal cultures treated with 3 μM Aβ show a sequential pattern of induction for components of the JAK-STAT pathway. 4h after Aβ exposure Tyk2 is upregulated in rat primary neuronal cultures. The levels of this gene are maintained elevated 24h after Aβ incubation. After Tyk2 levels are induced Jak1 and Jak2 are then subsequently upregulated.

We proceeded to test the response of the ak-StatJAK-STAT pathway in an acute in vivo rat model of Aβ toxicity (QPS; Austria GmbH). Five male rats were subjected to bilateral intracerebroventricular injection of 50 μM Aβ1-42 or equal volumes of phosphate buffered saline (PBS). 3h post injection, animals were sacrificed and brains harvested for RNA extraction. We measured the mRNA levels for Jak1, Jak2, Jak3 and Tyk2 in entorhinal cortex from Aβ-injected rats or PBS-sham controls by real time PCR. 3h after Aβ injection, the levels of Jak1 (p-value 0.001), Jak2 (0.0071) and Tyk3 (0.0032) were significantly upregulated (p<0.05) in the brains of these rats but not in control animals (Fig. 6) providing further validation for the implication of this pathway in Aβ-mediated toxicity.

**Fig. 6.**
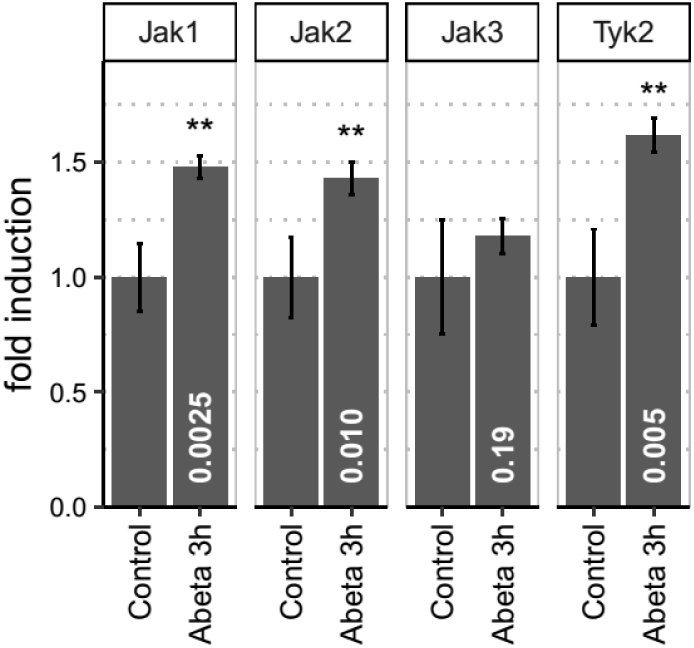
JAK kinases are upregulated in an in vivo model of Aβ toxicity Entorhinal cortex of male wild type rats injected with Aβ (n=5) or with PBS (n=5) bilaterally into the lateral ventricles was analyzed by real time PCR to determine the levels of Jak1, Jak2, Jak3 and Tyk2. Each sample was run in triplicate. Jak1, Jak2 and Tyk2 were significantly upregulated (p=0.001, p=0.0071, p=0.0032 respectively) in the Aβ-injected animals but not in the PBS injected control group. Data represented as normalized fold induction. Values given as mean ±s.e.m. Statistical significance determined by Student´s t-test.

## Discussion

We have presented here a series of integrated analyses predicated on the underlying hypothesis that co-morbidity of disease can, in some cases, indicate shared genetic susceptibility to disease and that this is manifested most robustly at the level of pathways more than at the level of single genes. By combining genome wide pathway association from all diseases together with their comorbidity with Alzheimer’s disease (AD), we identify JAK-STAT signaling as a shared factor correlated with the degree of comorbidity. In the first data-driven phase utilizing gene association data we find this disease cluster to include a series of disorders of immunity and inflammation together with Age Related Macular Degeneration (ARMD) and Type 1 Diabetes Mellitus (T1DM). The association with ARMD is particularly interesting as it has previously been found to be a risk factor for AD *(16, 17)*, because Aβ is a component of the drusen pathology in the retina of people with ARMD *(18, 19)* and because the gene most associated with ARMD – CFH - encodes for a protein replicated as a biomarker of AD, Complement Factor H *(20–22)*. That our approach of using all-GWAS data in a pathway clustering analysis identifies a relationship between AD and ARMD is all the more remarkable in because this association, in our study, is not driven by CFH. The association we find with T1DM is also intriguing as, although T2DM diabetes is one of the most substantiated risk factors for AD *(23)*, our data now suggests that there might be a relationship also with early onset T1DM that is worth further attention.

However, the most extensive association between shared pathways and disease we find is with disorders of immunity and inflammation. The role of inflammation in AD has been apparent for many years. This evidence is very extensive and comes from many directions *(24)*. It includes evidence the use of non-steroidal anti-inflammatory drugs appear to decrease the risk of AD *(25–27)*, post-mortem studies showing that inflammation is associated with AD pathology *(28, 29)*, and in vivo data showing that markers of inflammation are predominant amongst protein biomarkers both in AD and in pre-dementia conditions *(30, 31)*. In addition, there is increasing evidence from GWAS studies and from rare mutations, that genes encoding proteins involved in immunity are amongst the most consistently associated with disease *(24)*. However, when considered in isolation, the pathways and processes identified by AD genetic studies are predominantly those of complement signaling and microglial function *(3–32, 33)*. These pathways are clearly important in disease with very considerable evidence to support their role, but the approach we have used here, triangulating between GWAS, real-world and empirical data, and including all disease and all genes, nominates a pathway as part of the aetiopathogenic process that is not identified by such AD gene focused studies.

As in any ‘Big Data’ approach, there are limitations both to the datasets available and to our use of them. First, in using the GWAS catalogue as a primary data-source, we limit ourselves to disease-gene associations where a significant number of genes have been identified. We do this in order to provide sufficient power for analysis, but acknowledge that the limit of 25 susceptibility genes to enable a disease to enter analysis is both arbitrary and dependent on the size and numbers of studies that happen to have been conducted to date. Almost certainly, we miss information as a consequence of data limitation. Secondly, by segregating genes into pathways we attempt to overcome the intrinsic limitation of GWAS studies, in that biology is mechanistically enacted at the level of pathway and not gene, let alone SNP. Given the large number of SNPs and genes in the human genome, two diseases may have no elements measured in GWAS, or even sequencing studies, in common and yet share an overlapping disease pathogenesis. Measuring association not with SNP but with multiple SNPs across a gene (‘gene-wide association’) is one attempt to overcome the limitation; here we go one step beyond this with a pathway-wide association approach. However, in attempting to derive such information from the GWAS data, we are severely limited by current understanding of biological pathways, which is rudimentary at best. This limitation is bound to hinder our derivation of knowledge from information in this context. Thirdly, in seeking to identify diseases comorbid with AD, we have crudely utilised a dataset of concurrent diagnoses, taking no account of some of the confounds or other concerns of conventional epidemiology. Indeed we cannot be sure whether the co-morbidity we observe is due to the disease itself or the drugs used to treat the disease. However, we note that similar claims-level analyses of real-world clinical data have recently proven valuable in studying genetic and environmental factors shared amongst diseases *(34)*

We accept the limitations of our approach described above. However, in mitigation of these potential limitations, the datasets we examine are huge; namely, all genetic studies with all genes and all diseases in the first phase, and a dataset of over 6 million people in the second.

Furthermore, we would suggest that some confounds are less critical in the analysis we present here. For example, in studies of risk and protection then clearly understanding direction of effect – whether it is the disease or the treatment that affects risk – is fundamental. However, it becomes less important, potentially irrelevant, where we are determining simply whether the same processes are involved, as both disease and treatment will at some level and in some cases affect the same molecular pathways, which are the axis of our analysis. Finally, despite the limitations of deriving knowledge from data using this approach, the fact that the findings replicate in observational molecular studies in man and in experimental studies in rodents offers strong support to the results.

The JAK-STAT signaling pathway, nominated as a potential target for therapy through data-driven genomics and real-world data in this study, is a key regulator of the response to mediators of inflammation, including cytokines, chemokines *(35)* and microglia activation *(36)*. Binding of cytokines (interleukin, interferon and growth factors) and other ligands (such as hormones) to their receptors increases tyrosine kinase activity of Janus Kinases (JAKs, including TYK2), which in turn phosphorylate the receptors and recruit STATs which are themselves then phosphorylated. Subsequent dimerization leads to nuclear translocation and transcription factor activity. Given this critical role in the modulation of the inflammatory response, it is not surprising that JAK-STAT signaling has previously been associated with inflammatory disease and targeted for therapeutics, some of which have been approved for clinical use *(37)*. The pathway has also been identified as of importance in relation to diabetes *(38)* but less often in relation to AD. Very high concentrations of Aβ have been shown to increase tyrosine phosphorylation and transcriptional activity in a Tyk2 dependent manner in rodent models, and tyrosine phosphorylation of STAT3 is increased in AD brain *(39)*, while inhibition of JAK-STAT3 signaling inhibits activation of astrocytes and microglia in animal models of neurodegeneration *(40)*. JAK-STAT signaling has also been identified as a component of plasticity, specifically Long Term Depression (LTD) *(41, 42)*, interesting not least because LTD and Long Term Potentiation (LTP) are regulated, in opposite directions, by GSK3, the predominant tau-kinase implicated in AD pathogenesis *(43–45)*.

In summary, a combined, sequential analysis of GWAS data agnostic to disease type, combined with real-world data of co-incidence of AD with other diseases, nominated JAK-STAT signaling amongst other pathways as a possible underlying pathogenetic mechanism shared across multiple diseases. Remarkably, these diseases – inflammatory disorders, ARMD and Diabetes had previously been implicated as risk factors for AD. Adding to the weight of evidence for JAK-STAT signaling in AD, we subsequently found altered gene expression of the pathway in multiple human and rodent datasets and in empirical studies of Aβ exposure in rodents, both in vitro and in vivo. These data are not the first to nominate JAK-STAT signaling for therapeutic intervention *(46)*, as experiments suggest that humanin (HN) and colivelin (CLN) protect against AD-related neurotoxicity thought its activity in JAK-STAT. However the combination of genetic, human and rodent, observational and empirical data, make a strong case to pursue this pathway as a target for therapies for AD, not least because clinically approved compounds already exist. This further suggests that this novel integration of orthogonal data is a promising approach to find novel targets in complex disorders.

## Materials and Methods

### Subhead 1: Overlap of susceptibility genes across human disease

In order to identify biological pathways shared across different diseases, we utilised the GWAS-catalogue *(5)* to obtain a list of all known gene associations with disease derived from GWAS studies. We used the Experimental Factor Ontology (EFO) *(47)* to identify disease studies, filtering by diseases with at least 25 associated genes, and only including the 25 strongest associated genes ranked by p-value where more than 25 genes have been found to show some association. For any two genes ‘α’ and ‘β’ sampled in the GWAS-catalogue, we then calculated the number of KEGG pathways that these 2 genes share, obtaining a gene × gene matrix of which we show a section in Fig. 1. In order to determine whether susceptibility genes of any given disease share more pathways with the susceptibility genes of any other given disease than expected from chance alone, we used a non-parametric Wilcoxon rank-sum test.

Formally, if we denote the gene × gene matrix as ‘n(α,β)’ (with ‘α’ and ‘β’ being any gene sampled in the GWAS-catalogue), ‘A’ as the set of genes associated with disease ‘A’, and ‘B’ as the set of genes associated with disease ‘B’, the first sample ‘S+’ of our test becomes:

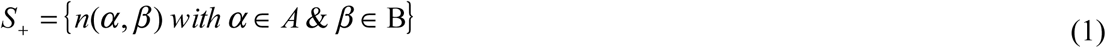

While the second sample ‘S-’ becomes:

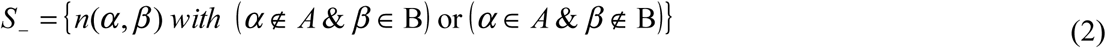

Then, a one-tailed Wilcoxon rank-sum test compares ‘S+’ against sample ‘S-’. The statistical results of this comparison determine whether diseases ‘A’ and ‘B’ share more pathways than expected from chance according to their GWAS associations.

### Subhead 2: Contribution of biological pathways to diseases

Given that the results from the analysis described above suggested an intersection between AD and inflammatory diseases and hence confirming known associations, as discussed in the results and discussion sections, we subsequently focussed on this overlap. For each inflammatory disease sampled in the GWAS-catalogue that had statistical power for subsequent analysis (see below), we calculated a so-called pathway load for each KEGG pathway. This is a numeric value that represents the proportion of susceptibility genes that a given disease has on a given KEGG pathway. Given disease ‘A’ and pathway ‘p’, the pathway load is equal to the number of susceptibility genes that disease ‘A’ has on pathway ‘p’, divided by the total number of times that any associated gene of disease ‘A’ is annotated as belonging to any KEGG pathway.

Formally, if we denote ‘m(A,p)’ as the number of genes of disease ‘A’ that belong to pathway ‘p’, the pathway load is:

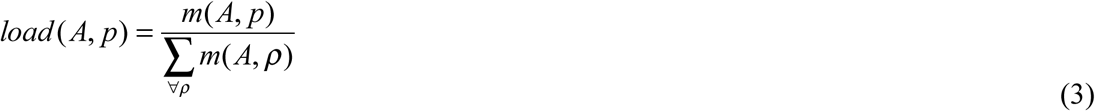

We divide by 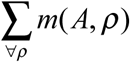 to control for diseases whose genes may have been more thoroughly studied, and therefore included in more KEGG pathways, than less studied diseases.

### Subhead 3: Shared contribution of biological pathways to disease comorbidity with AD

We determine the comorbidity of inflammatory diseases with AD using the US National Hospital Discharge Survey (NHDS https://www.cdc.gov/nchs/nhds/, *(48)*) records from 1979 to 2006, which can be found in the Inter-university Consortium for Political and Social Research (ICPSR), file ICPSR 24281 (http://www.icpsr.umich.edu/icpsrweb/ICPSR/studies/24281), or in the National Archive of Computerized Data on Aging (NACDA *(49)*). This randomised survey collects medical information, including final diagnosis coded in ICD-9-CM and demographic data, from the discharge records of a number of hospitals in the US, totalling 6,552,504 patient discharges in the sample used here. The record only includes hospitals with an average patient stay length of 30 days or less, excluding military and hospitals from institutions such as prisons *(48)*. We identified all patients in the dataset with a diagnosis of AD (ICD-9-CM code 331.0) and aged above 60 years in order to focus on late onset AD cases. For each case we find one control matched in gender, age, race and survey year (i.e. the year at which the survey was ran on that patient). To eliminate the diagnoses with lowest statistical power, we analyse only the inflammatory diseases that had more than 15 patients among the AD cases and their matched controls. For each inflammatory disease, we calculate the number of patients of that disease in the AD population divided by the number of patients of that disease in the matched population. For descriptive purposes we call this value ratioAD(A). Finally, for each KEGG pathway, we calculate whether the pathway load (load(A,p)) values correlate with the AD ratios (ratioAD(A)) using Pearson’s correlation.

### Subhead 4: Pathway gene expression dysregulation in AD

In order to examine the relationship between gene expression and disease we obtained RNA expression data from 4 independent datasets; 1) AddNeuroMed (ANM), a longitudinal multi-centre cohort study with blood samples from 105 AD cases, 125 Mild Cognitive Impairment (MCI) individuals and 114 controls *(6, 50)*; 2) the Dementia Case Register (DCR), a single centre longitudinal cohort utilising the exact same protocol as ANM with blood samples from 90 AD cases, 65 MCI individuals and 73 controls; 3) the largest post-mortem AD dataset available in ArrayExpress, E-GEOD-44770, with samples of post-mortem pre-frontal cortex from 129 AD cases and 101 controls *(7)*; 4) a rodent in-vitro dataset of neuronal gene response to amyloid peptide *(8)*. Throughout this paper, these datasets are respectively denoted “Dataset 1”, “Dataset 2”, “Dataset 3” and “Dataset 4”. The case ascertainment, diagnostic process and other details of ANM have been described previously *(6–50, 51)* while DCR used an identical clinical and laboratory protocol. Demographics from the three human cohorts (Datasets 1 to 3) are reported in table 2.

**Table 2.** Characteristics of human datasets. Human study cohorts with the variables used in the linear model (ANM-AddNeuroMed; DCR-dementia case register; AE-ArrayExpress – see methods for details)

|  | Dataset 1 | Dataset 2 | Dataset 3 |
| --- | --- | --- | --- |
|  | (blood, ANM) | (blood, DCR) | (brain, AE) |
| Gender (n) |  |  |  |
| Female | 209 | 148 | 108 |
| Male | 135 | 80 | 121 |
| Age (years) |  |  |  |
| mean ± SD | 75.9 ± 6.8 | 78.1 ± 7.0 | 73.8 ± 12 |
| Centre (n) |  |  |  |
| 1 | 44 | 44 | 229 |
| 2 | 34 | 11 | - |
| 3 | 84 | 62 | - |
| 4 | 43 | 43 | - |
| 5 | 45 | 25 | - |
| 6 | 94 | 43 | - |
| Diagnosis (n) |  |  |  |
| AD | 105 | 90 | 129 |
| control | 114 | 73 | 100 |
| MCI | 125 | 65 | 0 |

To test for statistically significant differences in RNA-expression in these 4 datasets for a given pathway, we use a per gene general linear model (GLM) with a binomial link function, which models AD status (two levels per person – either AD patient or control) as a function of RNA-expression while controlling for a number of covariates (gender, age and sampling centre). The p-values we report correspond to the RNA-expression variable per gene.

### Subhead 5: Proof of concept in an in vitro rat model of Aβ exposure

In order to explore JAK-STAT signaling in vitro, primary neuronal cultures were generated from Sprague Dawley E18 rat embryos by papain dissociation according to the manufacturer’s instructions (Worthington, Lakewood, NJ, USA) and cultured as previously described *(52)*. Briefly, brains were harvested and maintained in sterile PBS, hippocampi and cortices were dissected out using a dissection microscope, triturated using a sterile glass Pasteur pipette and maintained in serum-free medium. Viable cells were counted using a hemacytometer. Neurons were plated in Neurobasal medium supplemented with B27, 0.30% glutamine, 100 U/ml penicillin and 100 μg/ml streptomycin, at a density of 300,000 cells/ml in plates coated with poly-D-lysine and incubated at 37°C in 5% CO2 atmosphere. Neuronal cultures were treated 7-9 days post-plating.

Aβ1–42 peptide was purchased from Dr. David Teplow (California, UCLA) and was resuspended in 100% 1,1,1,3,3,3 hexafluoro-2-propanol (HFIP) at a final concentration of 1 mM. For complete solubilisation the peptide was homogenized using a Teflon plugged 250 μl Hamilton syringe. HFIP was removed by evaporation in a SpeedVac, Aβ1–42 resuspended at a concentration of 5 mM in dimethylsulfoxide (DMSO) and sonicated for 10 minutes. Oligomers were prepared as previously described *(53)*: Aβ1–42 was diluted in PBS to 400 μM and 1/10 volume 2% sodium dodecyl sulfate (SDS) in H2O added. Aβ was incubated for 24 hours at 37°C and further diluted to 100 μM in PBS followed by 18 hours incubation at 37°C. Rat primary neuronal cultures were treated with 3 μM Aβ for either 4h or 24h. These time points were selected based on the commonly observed time progression of changes in RNA after Aβ treatment.

### Subhead 6: Proof of concept in an in vivo rat model of Aβ exposure

We further tested whether the observations made in vitro were also found in in vivo models. Male wild type Wistar rats (approximately 300g body weight) were subjected to bilateral injections of 50 μM Aβ1-42 or PBS (n = 5 per group) into the lateral ventricles (QPS, Austria). Brains were collected 3 hours post injection. Entorhinal cortex was subdissected, frozen in liquid nitrogen and processed for RNA extraction. Frozen tissues were thawed and homogenised in Trizol and total RNA extracted according to the manufacturer’s instructions (Invitrogen, Paisley, UK).

Total RNA from entorhinal cortex (1 μg) was reverse transcribed using random hexamers and a Taqman RT kit (Applied Biosystems, Cheshire, UK) according to the manufacturer’s instructions. PCR primers were designed using the Universal Probe Library package (Roche Molecular Biochemicals, Lewes, UK) and used in SYBR Green-based PCR reactions performed on a StepOnePlusTM (96 well) thermal cycler (Applied Biosystems, California, USA). Relative quantification of gene expression between samples was determined using the 2^-ΔΔCT^ method as described by Livak and Schmittgen *(54)*. Internal control genes used to normalise for RNA input were HPRT and β-actin.

Primer sequences: Jak1(NM_053466.1): Forward: 5´-ccaccgggacatttcact-3´; Reverse: 5´-ttgtgggaaacctgtctcatc-3´; Jak2 (NM_031514.1): Forward: 5´-ggagagtatgttgccgaagaa-3´; Reverse: 3´-atattatgatacacaggcgtaatacca-3´; Jak3 (NM_012855.2): Forward: 5´-ggccaaagtcccatcttct-3´; Reverse: 5´-gaagctccacacgtcagattg-3´; Tyk2 (NM_001257347.1): Forward: 5´-tgccatcttgctctcaacc-3´; Reverse: 5´-gtgagggatacagttcttgaagc-3´.

All samples were run in triplicate from three independent experiments. The mean crossing threshold (CT) values for both the target and internal control genes in each sample were determined and the 2^-ΔΔCT^ calculations performed. The fold change in the target genes (after normalising to the internal control gene) were calculated for each sample and the mean calculated. Statistical significance was determined by Student´s t-test. Data are represented as normalised fold increases over control samples. Values are given as mean ± s.e.m (standard error of the mean).

### Subhead 7: Ethical considerations

All animal studies described in this manuscript were ethically reviewed and carried out in accordance with Animals (Scientific Procedures) Act 1986. We further certify that the research (including Embryonic and Fetal Research if appropriate) was conducted according to the requirements of POL-GSKF-410 and associated relevant SOPs, and that all related documentation is stored in an approved HBSM database. Human biological samples were sourced ethically and their research use was in accord with the terms of the informed consents

## Supplementary Materials

Supplementary materials are detailed in “Supplementary materials.doc” file, as requested in the instructions for authors provided at “http://stm.sciencemag.org/content/instructions-authors-new-research-articles”.

Fig. S1. Overlap of GWAS genes, full version.

## Acknowledgments

The work described in this manuscript is part of the Med Bioinformatics consortium (http://www.medbioinformatics.eu/), and we thank its inestimable help and assistance during the development of this study.

## Funding

This work was supported by the Wellcome Trust [104025] and the European Union’s Horizon 2020 research and innovation programme 2014-2020 under Grant Agreement No 634143.

## Author contributions

Here describe the contributions of each author (use initials) to the paper.

The study was conceived and designed by AJN-H and SL and data analysis conducted by AJN-H, LF and MA-M. Experimental studies in vitro and in vivo were conducted by ER and LT and the manuscript edited by AJNH, JQ, JDR, JC and SL. Members of the NIMA consortium are listed below:

NIMA – Wellcome Trust Consortium for Neuroimmunology of Mood Disorders and Alzheimer’s Disease

Consortium members

Cambridge

Edward T. Bullmore (PI, EC)^1,2,11^, Junaid Bhatti^1^, Samuel J. Chamberlain^1,2^, Marta M. Correia^1,12^, Amber Dickinson*, Andy Foster^2^, Manfred Kitzbichler^1^, Clare Knight^2^, Mary-Ellen Lynall^1^, Christina Maurice^1^, Howard Mount^13^, Ciara O'Donnell^1^, Linda J. Pointon^1^, Peter St George Hyslop^1,13,14^, Lorinda Turner^1^, Barry Widmer^1^, Guy B. Williams^1,14^

Cardiff

B. Paul Morgan (PI)^15^, Claire Leckey^15^, Angharad Morgan^15^, Caroline O'Hagan*, Samuel Touchard^15^

Glasgow

Jonathan Cavanagh (PI, EC)^3^, Catherine Deith*, John McClean^16^, Alison McColl^3^, Andrew McPherson*, Paul Scouller*, Murray Sutherland^16^

Independent advisor

H.W.G.M. (Erik) Boddeke (EC)^17^

GSK

Jill Richardson (EC)^18^, Shahid Khan^11^, Phil Murphy^19^, Christine Parker^19^, Jai Patel^11^

Janssen

Declan Jones (EC)^6^, Peter de Boer^4^, John Kemp^4^, Paul Acton^6^, Wayne C. Drevets^6^, Jeffrey S. Nye (deceased), Gayle Wittenberg^6^, John Isaac^6^, Anindya Bhattacharya^6^, Nick Carruthers^6^, Hartmuth Kolb^6^

Kings College London

Carmine Pariante (PI)^10^, Gareth Barker^20^, Heidi Byrom^10^, Diana Cash^20^, Antony Gee^20^, Caitlin Hastings^10^, Nicole Mariani^10^, Anna McLaughlin^10^, Valeria Mondelli^10^, Maria Nettis^10^, Naghmeh Nikkheslat^10^, Karen Randall^20^, Hannah Sheridan*, Camilla Simmons^20^, Nisha Singh^20^, Federico Turkheimer^20^, Victoria Van Loo*, Marta Vicente Rodriguez^20^, Tobias Wood^20^, Courtney Worrell*, Zuzanna Zajkowska*

Lundbeck

Niels Plath (EC)^21^, Jan Egebjerg^21^, Hans Eriksson^21^, Francois Gastambide^21^, Karen Husted Adams^21^, Ross Jeggo^21^, Christian Thomsen^21^, Jason O'Connor^22^, Jan Torleif Pederson^21^, Brian Campbell*, Thomas Möller*, Bob Nelson*, Stevin Zorn*

Oxford

Mary Jane Attenburrow (PI)^7,23^, Alison Baird, Jithen Benjamin^23^, Stuart Clare^25^, Philip Cowen^7^, I-Shu (Dante) Huang^24^, Samuel Hurley*, Helen Jones^23^, Simon Lovestone^7^, Francesca Mada^23^, Alejo Nevado-Holgado^7^, Akintayo Oladejo*, Elena Ribe^7^, Anviti Vyas*

Pfizer

Zoe Hughes (EC)^26^, Rita Balice-Gordon*, Brendon Binneman^26^, James Duerr^26^, Terence Fullerton^26^, Justin Piro^26^, Tarek Samad^26^, Jonathan Sporn^26^

Southampton

Hugh Perry (PI)^27^, Madeleine Cleal*, Gemma Fryatt^27^, Diego Gomez-Nicola^27^, Renzo Mancuso^27^

Sussex

Neil Harrison (PI, EC)^28^, Mara Cercignani^28^, Charlotte Clarke^28^, Elizabeth Hoskins^29^, Charmaine Kohn^29^, Rosemary Murray*, Dominika Wlazly^30^

PI = Principal Investigator

EC = Executive committee member

**Competing interests**:

Jill Richardson is an employee of GSK, and holds GSK stocks and shares.

Jonathan Cavanagh holds a Wellcome Trust strategic award which involves industrial-academic collaboration with Janssen & Janssen, GSK, Pfizer and Lundbeck.

Jonathan Cavanagh has been on a scientific advisory board for GSK.

Simon Lovestone has provided consultancy services to SomaLogic, Eisai, OptumLabs and J&J and is in receipt of direct grant funding from J&J and Astra Zeneca

## Footnotes

1 Department of Psychiatry, School of Clinical Medicine, University of Cambridge, CB2 0SZ, UK

2 Cambridgeshire and Peterborough NHS Foundation Trust, Cambridge, CB21 5EF, UK

3 Sackler Centre, Institute of Health & Wellbeing, University of Glasgow, Sir Graeme Davies Building, Glasgow, G12 8TA, UK

4 Neuroscience, Janssen Research & Development, Janssen Pharmaceutica NV, Turnhoutseweg 30, B-2340, Beerse, Belgium

5 The Maurice Wohl Clinical Neuroscience Institute, Cutcombe Road, London, SE5 9RT, UK

6 Neuroscience, Janssen Research & Development, LLC, Titusville, NJ, 08560, USA

7 University Department of Psychiatry, Warneford Hospital, Oxford, OX3 7JX, UK

8 Brighton & Sussex Medical School, University of Sussex, Brighton, BN1 9RR, UK

9 Sussex Partnership NHS Foundation Trust, Swandean, BN13 3EP, UK

10 Stress, Psychiatry and Immunology Lab & Perinatal Psychiatry, Maurice Wohl Clinical Neuroscience Institute, Kings College, London, SE5 9RT, UK

11 Immuno-Psychiatry, Immuno-Inflammation Therapeutic Area Unit, GlaxoSmithKline R&D, Stevenage SG1 2NY, UK

12 MRC Cognition and Brain Sciences Unit, 15 Chaucer Road, Cambridge CB2 7EF, UK

13 Tanz Centre for Research in Neurodegenerative Diseases, 60 Leonard Avenue, Toronto, ON M5T 2S8 Canada

14 Department of Clinical Neurosciences, University of Cambridge, CB2 0SZ, UK

15 University of Cardiff, Cardiff CF10 3AT, UK

16 NHS Greater Glasgow and Clyde, 1055 Great Western Rd, Glasgow G12 0XH, UK

17 University of Groningen, 9712 CP Groningen, Netherlands

18 Neurosciences Virtual PoC DPU, GlaxoSmithKline R&D, Stevenage SG1 2NY, UK

19 Experimental Medicine Imaging, GlaxoSmithKline R&D, Stevenage SG1 2NY, UK

20 Centre for Neuroimaging Sciences, Denmark Hill, London SE5 9AF, UK

21 H. Lundbeck A/S Ottiliavej 9, 2500, Valby, Denmark

22 University of Texas Health Science Center at San Antonio, 7703 Floyd Curl Dr, San Antonio, TX 78229, USA

23 NIHR Oxford cognitive health Clinical Research Facility, Warneford Hospital, Oxford, OX3 7JX, UK

24 The Kennedy Institute of Rheumatology, Roosevelt Dr, Oxford OX3 7FY, UK

25 Oxford Centre for Functional MRI of the Brain, John Radcliffe Hospital, Oxford OX3 9DU, UK

26 Pfizer, Inc, 1 Portland Street, Cambridge MA, USA

27 Centre for Biological Sciences, University of Southampton, Southampton, UK

28 Clinical Imaging Sciences Centre (CISC), University of Sussex, Brighton, BN1 9RR, UK

29 Sussex Partnership NHS Foundation Trust, Nevill Avenue, Hove BN3 7HZ, UK

30 Brighton & Sussex University Hospitals NHS Trust, Brighton BN2 5BE, UK

* Former consortium members

